# Interpretable modeling of time-resolved single-cell gene-protein expression using CrossmodalNet

**DOI:** 10.1101/2023.05.16.541011

**Authors:** Yongjian Yang, Yu-Te Lin, Guanxun Li, Yan Zhong, Qian Xu, James J. Cai

## Abstract

Cell-surface proteins play a critical role in cell function and are primary targets for therapeutics. CITE-seq is a single-cell technique that enables simultaneous measurement of gene and surface protein expression. It is powerful but costly and technically challenging. Computational methods have been developed to predict surface protein expression using gene expression information such as from single-cell RNA sequencing (scRNA-seq) data. Existing methods however are computationally demanding and lack the interpretability to reveal underlying biological processes. We propose CrossmodalNet, an interpretable machine learning model, to predict surface protein expression from scRNA-seq data. Our model with a customized adaptive loss accurately predicts surface protein abundances. When samples from multiple time points are given, our model encodes temporal information into an easy-to-interpret time embedding to make prediction in a time point-specific manner able to uncover noise-free causal gene-protein relationships. Using two publicly available time-resolved CITE-seq data sets, we validate the performance of our model by comparing it to benchmarking methods and evaluate its interpretability. Together, we show our method accurately and interpretably profiles surface protein expression using scRNA-seq data, thereby expanding the capacity of CITE-seq experiments for investigating molecular mechanisms involving surface proteins.

## 1. Introduction

Single-cell RNA sequencing (scRNA-seq), which allows transcriptomic data collection from thousands of cells in parallel (Quake, 2021), enables examination of cellular states at individual cell level, leading to insights into diverse cell type identification, gene regulation, and cellular communication (Jindal et al., 2018; Osorio et al., 2020; 2022; Yang et al., 2023). Compared to traditional single-cell techniques that measure only one aspect of cellular activity, the ability of multimodal (Ling et al., 2023; Han et al., 2022a;b) approaches has the potential to significantly improve our understanding of cellular behavior and function, thereby shedding light on a vast array of biological questions. Cellular Indexing of Transcriptomes and Epitopes by Sequencing (CITE-seq) (Stoeckius et al., 2017) is a cutting-edge sequencing method that allows simultaneous measurement of gene and surface protein expression at the single-cell level. CITE-seq, however, faces some challenges. First, CITE-seq experiments are costly and require specialized equipment and trained personnel. Second, the number of available antibodies limits the number of surface proteins that CITE-seq can measure. This is problematic when attempting to analyze complex cell populations. Antibody cross-reactivity and non-specific binding may also result in false CITE-seq discoveries (Restani et al., 2002).

Machine learning methods have been developed to learn the relationship between genes and proteins and translate between single-cell measurements of these two modalities. Seurat 4 (Hao et al., 2021) and totalVI (Gayoso et al., 2021), for instance, have been developed, but their computational cost is especially high. Recent work scIPENN (Lakkis et al., 2022) is a multi-use framework for CITE-seq and scRNA-seq integration with surface protein prediction and imputation. However, scIPENN’s RNN blocks could cause the gradient vanishing problem and potentially hinder the training process (Pascanu et al., 2013). More importantly, the inability to interpret what and how genes significantly regulate protein expression levels over time may limit its application in understanding essential cellular system questions.

Here we propose CrossmodalNet, an interpretable machine learning model with customized adaptive loss that learns to translate between modalities of genes and proteins using CITE-seq data while encoding temporal information, and can accurately predict protein expression using only scRNA-seq data at chosen time points. We evaluate the performance of our model using two publicly available CITE-seq data sets containing hematopoietic stem and progenitor cells (HSPCs) and peripheral blood mononuclear cells under development, respectively. By combining the interpretability of linear models with the flexibility of non-linear models, we show that our model decomposes transcriptional information of cells into basal and temporal domain, with the latter forming an easy-to-interpret time embedding. Using the learned time embedding, we demonstrate that our model outperforms other benchmarking methods for protein prediction at both observed and unobserved time points. Moreover, we show our model is capable of elucidating noise-free causal gene-protein relationships that are typically investigated in large-scale genomic studies.

## 2. Methods

### 2.1. Notations

Let *X ∈* ℝ^*m×g*^ and *Y ∈* ℝ^*m×p*^ represent gene and protein expression profiles from CITE-seq, respectively, with *m* cells, *g* genes, and *p* proteins. We are given a training data set 𝒟 = (*x*_1_, *y*_1_, *t*_1_), …, (*x*_*m*_, *y*_*m*_, *t*_*m*_), and our objective is to learn *y*_*i*_ given *x*_*i*_ at discrete time *t*_*i*_ for each cell *i* using *𝒟*.

### 2.2. Proposed model

We recruit a framework similar to Fader Networks (Lample et al., 2017) that enables interpretable covariets in addition to nonlinear mapping between gene and protein expression and use multitask training strategy to optimize the model performance (Figure 1). Recent work CPA (Lotfollahi et al., 2023) and its further research MultiCPA (Inecik et al., 2022), for example, leverage the Fader Networks for predictions of drug responses and genetic perturbations. Details are introduced as follows.

**Figure 1.**
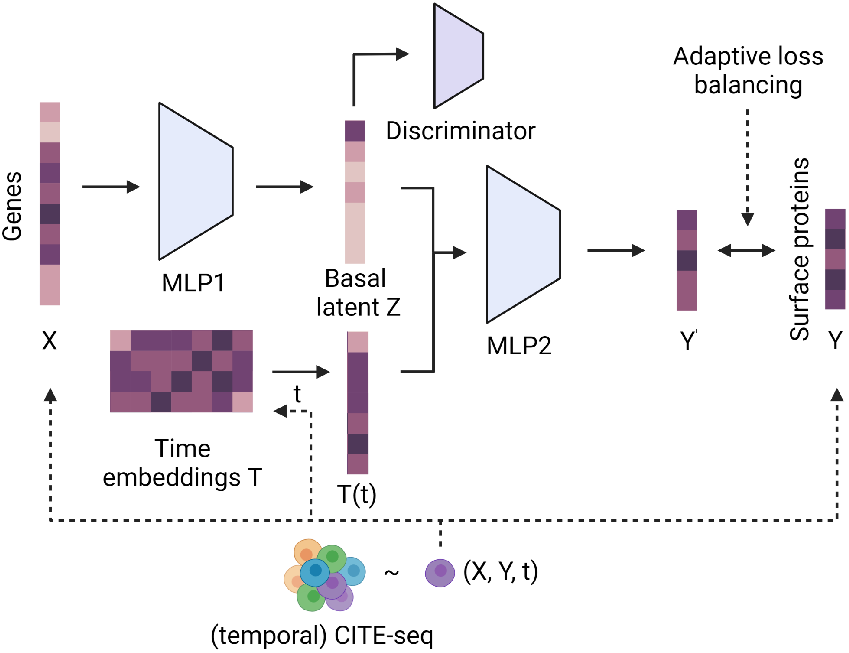
The framework of CrossmodalNet predicting protein expressions given gene expression profiles using temporal CITE-seq data.

### 2.3. Fader Networks

#### 2.3.1. Model architecture

We first initialize a learnable time embedding *T ∈* ℝ^*c×d*^, where *c* is the number of unique class of time and *d* represents the latent dimension of the model. We denote the *d*-dimensional time representation as *T*_*i*_ := *T* (*t*_*i*_). Let *C*_*k*_ be a Linear-BatchNorm block with *k* output features. The first MLP (MLP1) consists of *C*_*d*_ *– ReLU – Dropout – C*_*d*_ and the second MLP (MLP2) consists of *C*_*d*_ *– ReLU – C*_*out*_ without BatchNorm (Santurkar et al., 2018) in the last output layer.

#### 2.3.2. Discriminator objective

We introduce a discriminator that calculates the probability of a time point *t*_*i*_ given the input *x*_*i*_. The objective function of the discriminator is defined as

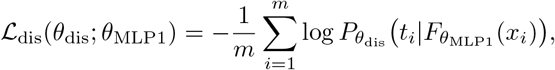

where *θ*_dis_ and *θ*_MLP1_ are parameters of discriminator and MLP1. A well-trained discriminator will enable a cell’s basal latent state disentangled from the time.

#### 2.3.3. Adversarial objective

Denote the the basal latent state as 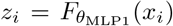, we next aggregate *z*_*i*_ of cell *i* and its time representation *T*_*i*_ into a unified space, and then map the sum to the protein expression. The Fader loss, given the discriminator parameters *θ*_dis_, is:

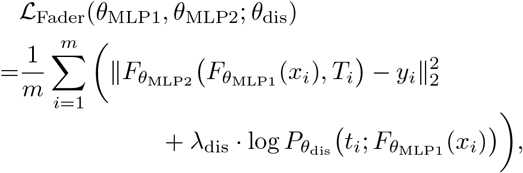

where *θ*_MLP2_ is parameters of MLP2 and *λ*_dis_ is a regularize parameter. Minimizing the training loss requires optimizing both the mean squared error (MSE) reconstruction loss, which is used to reconstruct *y*_*i*_, and the cross entropy loss, which is used to predict *t*_*i*_. We denote the reconstructed protein expression by

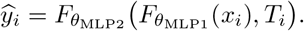

### 2.4. Multitask training

We introduce a new loss function called negative log-correlation (NLC) loss (Figure S1). The NLC loss directly regulates the correlation between the predicted values 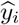 and the actual values *y*_*i*_, and can be used for backpropagation. The formula for the NLC loss is:

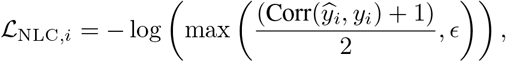

where *ϵ* is a small value to stable the computation. Denote the NLC loss by

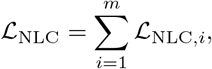

and the total loss of the entire model is defined as:

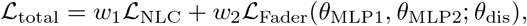

where *w*_1_ and *w*_2_ are adaptive parameters balancing the two losses.

We employ GradNorm method proposed by (Chen et al., 2018) to optimize the loss function, which would improve model performance and reduce overfitting when compared to single-task models. To this end, we define

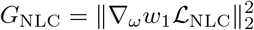

and

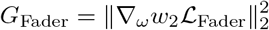

as the *L*_2_ norm of the gradient of the weighted single-task loss with respect *ω*, where we choose *ω* as the first layer parameters of *θ*_MLP1_. We also define the average gradient norm across all tasks as 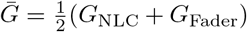. The goal of GradNorm is to match the scale of *G*_NLC_ and *G*_Fader_. Let ℒ_NLC_(*t*) and ℒ_Fader_(*t*) be the loss function values at the *t*-th iteration, we define

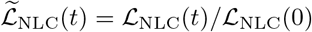

as the NLC loss ratios at time *t*, where ℒ_NLC_(0) is the loss at initialization, and 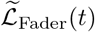 and ℒ_Fader_(0) are defined similarly. We also define the average loss function value at time *t* as 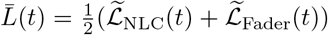, and the relative inverse training rate by 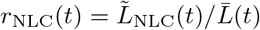 and *r*_Fader_(*t*) similarly. Algorithm 1 explicitly demonstrates the training with CrossmodalNet.

### 2.5. Hyperparameter tuning and implementation

We perform a random hyperparameter search using Ray Tune v2.0.0 (Liaw et al., 2018) of 100 trials. Table S1 outlines the distribution of values for hyperparameter search. To implement, we first split cells into three data sets for training (80%), validation (5%), and testing (15%). For in-distribution predictions, cells are randomly sampled such that proportion of cells at different time points remain equal in each set. For out-of-distribution predictions, cells at a given time point are kept as the testing set, while other cells are treated as the training and validation set. The maximum iteration number was set to 500, and early stopping is added after Pearson correlation coefficient of validation set reaches the maximum for 10 iterations. The Adam optimizer is used for all trainings.

#### Algorithm 1 Training with CrossmodalNet

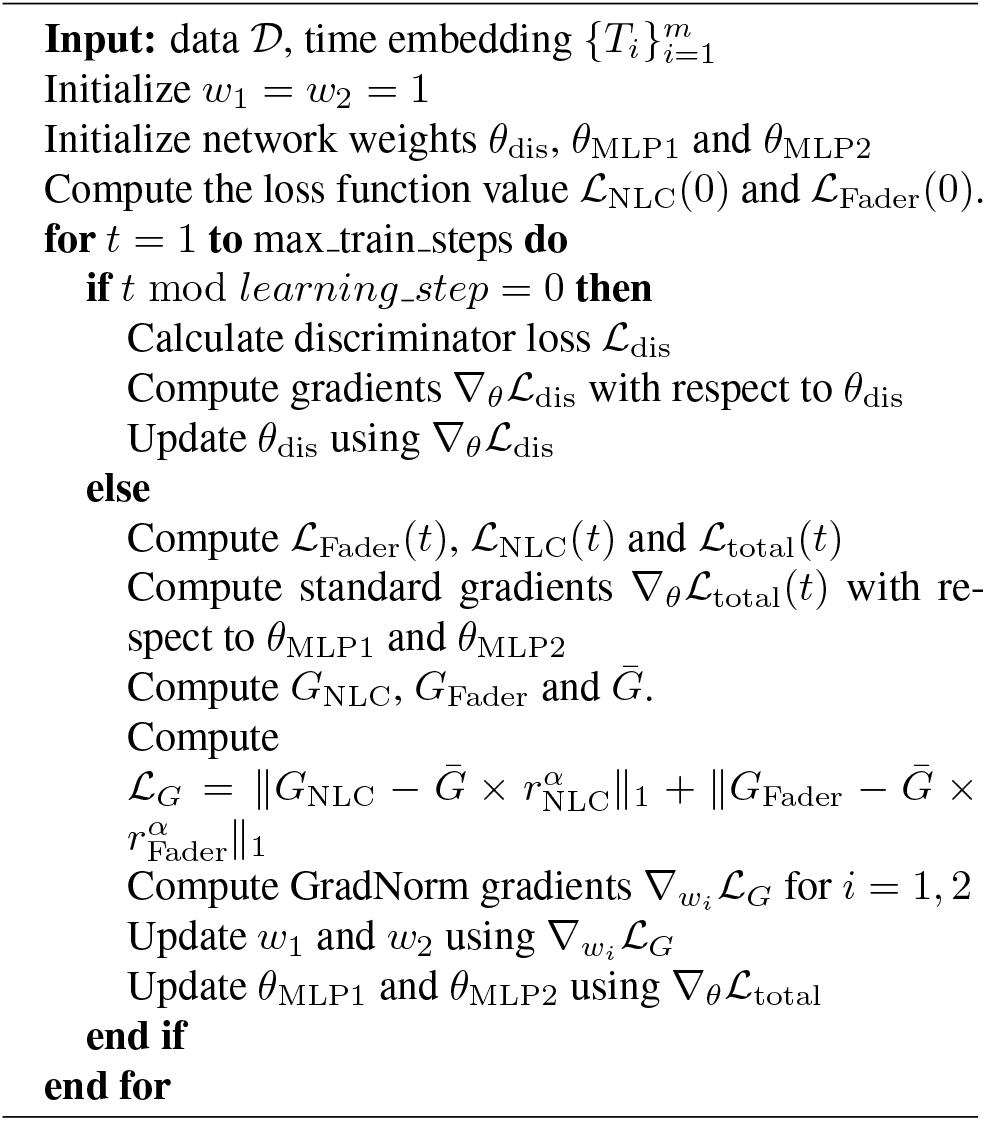

### 2.6. Causal gene-protein relationship inference

Given the nonlinearity of CrossmodalNet, we apply saliency maps (Simonyan et al., 2013) to differentiate the importance of input features for output. In the case of causal gene-protein relationship analysis, given a protein *j*, its saliency with respect to genes can be computed by aggregating all cells:

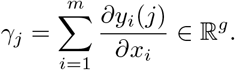

Since our latent inference only contains basal information, we anticipate the saliency analysis will reveal more noise-free gene-protein relationships.

## 3. Experiment setup

### 3.1. Data

We use two real CITE-seq data sets for model training and evaluation.

#### HSPC data set

This data set was collected from over 70,000 mobilized peripheral CD34+ HSPCs isolated from four healthy human donors across three time points from Kaggle Open Problems in Single-Cell Analysis (Cellarity et al., 2022) guided by (Velten et al., 2017) where 140 surface proteins were measured.

#### Myeloid data set

This data set was collected from over 47,000 peripheral blood mononuclear cells of patients with advanced biliary tract cancer (BTC) across three time points following anti-PD-1 treatment (Keenan et al., 2022) where 99 surface proteins were measured. Four CD14+ monocyte sub-populations by responsive BTC patients are used.

### 3.2. Preprocessing

We perform RNA library-size normalization and log1p transformation on scRNA-seq data using NormalizeData function from Seurat v4.0.2 package (Hao et al., 2021). We perform dsb transformation for surface protein data using DSBNormalizeProtein function from dsb package v1.0.2 (Mule` et al., 2022). Both normalization methods are performed after data split. All default settings are retained and used.

### 3.3. Model evaluation

We evaluate our model against several baseline methods including linear regression, ridge regression, lightGBM (Ke et al., 2017), MLP, and scIPENN. ScIPENN consequently provides more accurate results than totalVI and Seurat 4, so we have excluded them in our evaluation. We utilize scikit-learn v1.2 to build the linear and ridge regression models with default hyperparameters. The LightGBM model is built using LightGBM v3.3.5, and we use randomized cross-validation to select the optimal hyperparameters. Pytorch v2.0 (Paszke et al., 2019) and pytorch-lightning v2.0.2 (Falcon et al., 2019) are used to create the MLP models. Similar to how we tune our model, we utilize Ray to determine the best model structure and hyperparameters. We evaluate each model’s performance using MSE and Pearson correlation coefficient.

## 4. Results

### 4.1. CrossmodalNet accurately translates between gene-protein modalities

We first demonstrate the performance and functionality of CrossmodalNet using two publicly available CITE-seq data. These two data sets represent two application scenarios of CrossmodalNet—i.e., homogeneous and heterogeneous cell types developing across time. Table 1 compares the performance of CrossmodalNet and other methods under the in-distribution setting. Our result indicates that CrossmodalNet is capable of attaining the highest Pearson correlation coefficient of all methods. In addition, even though linear models show good predictive performance, they are not superior to lightGBM and MLPs. In particular, MLP trained with MSE loss achieves the lowest MSE, which is expected. It should be noted, however, that this MSE model does not produce the highest Pearson correlation coefficient, indicating that a single MSE loss may not be the optimal choice for this task.

**Table 1.** In-distribution predictions.

| DATA SET | HSPC |  | MYELOID |  |
| --- | --- | --- | --- | --- |
| METHOD | CORR | MSE | CORR | MSE |
| CROSSMODALNET | **0.593** | 0.259 | **0.819** | 0.249 |
| MLP+MSE | 0.561 | **0.140** | 0.792 | 0.160 |
| MLP+NLC | 0.490 | 0.316 | 0.560 | 0.670 |
| LIGHTGBM | 0.575 | 0.259 | 0.791 | **0.151** |
| LINEAR REG. | 0.559 | 0.269 | 0.745 | 0.301 |
| RIDGE REG. | 0.560 | 0.268 | 0.772 | 0.304 |
| SCIPENN | 0.379 | 0.456 | 0.429 | 0.489 |

### 4.2. CrossmodalNet generalizes to unobserved time point

To demonstrate the generalization of the CrossmodalNet model, we hold out cells at an intermediate time point and train with cells preceding and following the time point. Specifically, we hold out HSPCs from day 3 and train day 2 and 4; monocytes from week 2 and train week 1 and 3, respectively. After obtaining the learnable time embedding, the unseen time representation is inferred through a linear interpolation between two learned vectors of the time embedding. During testing, we manually concatenate this inferred time representation vector with basal latent representations of cells given by the trained model to obtain predictions. In table 2, we compare the performance of CrossmodalNet and other methods under this setting. Our results show that CrossmodalNet outperforms other methods, indicating its high generalizability. Interestingly, most models trained with myeloid data exhibit relatively inferior performance relative to in-distribution predictions, whereas models trained with HSPC data do not compromise. This might reflect a more prominent temporal batch effect in myeloid data, which cannot be modeled linearly. In addition, we see that the Pearson correlation coefficients of MLP models trained with NLC loss are greater than those with MSE loss. This observation suggests that our NLC loss may aid neural networks at a certain level of generalization.

**Table 2.**
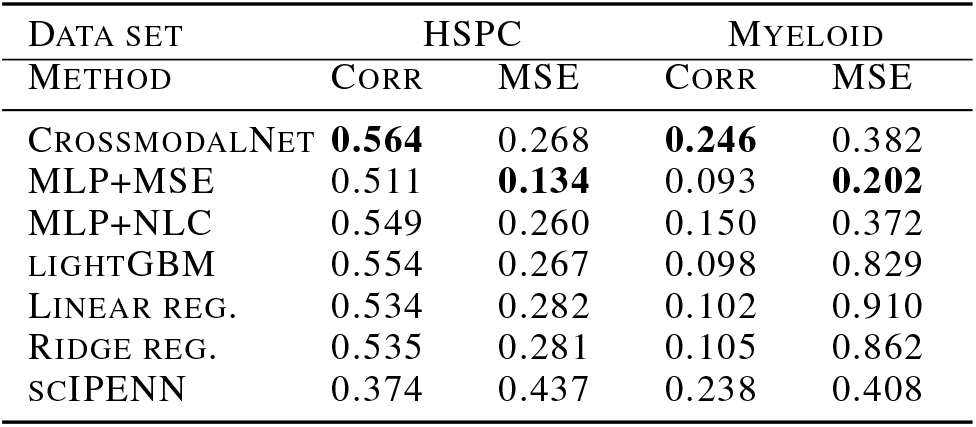
Out-distribution predictions.

### 4.3. CrossmodalNet infers causal gene-protein relationships

The development of HSPCs is tightly regulated by changes in gene and protein expression, but there is currently limited understanding of how these two measurements co-vary in HSPCs when they develop into more mature blood cells. The first CITE-seq data set contains more than 70,000 CD34+ HSPCs based in (Velten et al., 2017), which suggests that discrete cell populations are established only when differentiation has progressed to the level of restricted progenitors associated with the upregulation of surface protein CD38.

We first illustrate the correlation of expression between the surface protein CD38 and its coding gene is low (0.411, Figure 2a) whereas our model’s prediction significantly improved it (0.708, Figure 2b). We visualize the learned high dimensional time embedding in PCA where three time vectors are found to be almost evenly separated from each other (Figure S2a). We compute the saliency of features to determine which features (genes) our model pays attention to for predicting the expression level of CD38. Figure 2c depicts the saliency ranking of the top ten most prominent genes. Gene *CD38* is at the top of the list despite its low expression level, which causes it to be obscured (Figure 2d) by other highly expressed genes, indicating that our method successfully recognizes the gene *CD38*’s significant contribution to the protein CD38’s expression. Gene Set Enrichment Analysis (GSEA) with the KEGG pathway database using top 100 saliency genes ranks *Hematopoietic cell lineage* at the top (Figure 4a), which corresponds to the potential central role of CD38 in cell differentiation presented in the study of HSPCs (Velten et al., 2017). Together, our results show our model accurately model the intrinsic gene-protein relationships across time and shed light on the molecular mechanism underlying the functioning of CD38.

**Figure 2.**
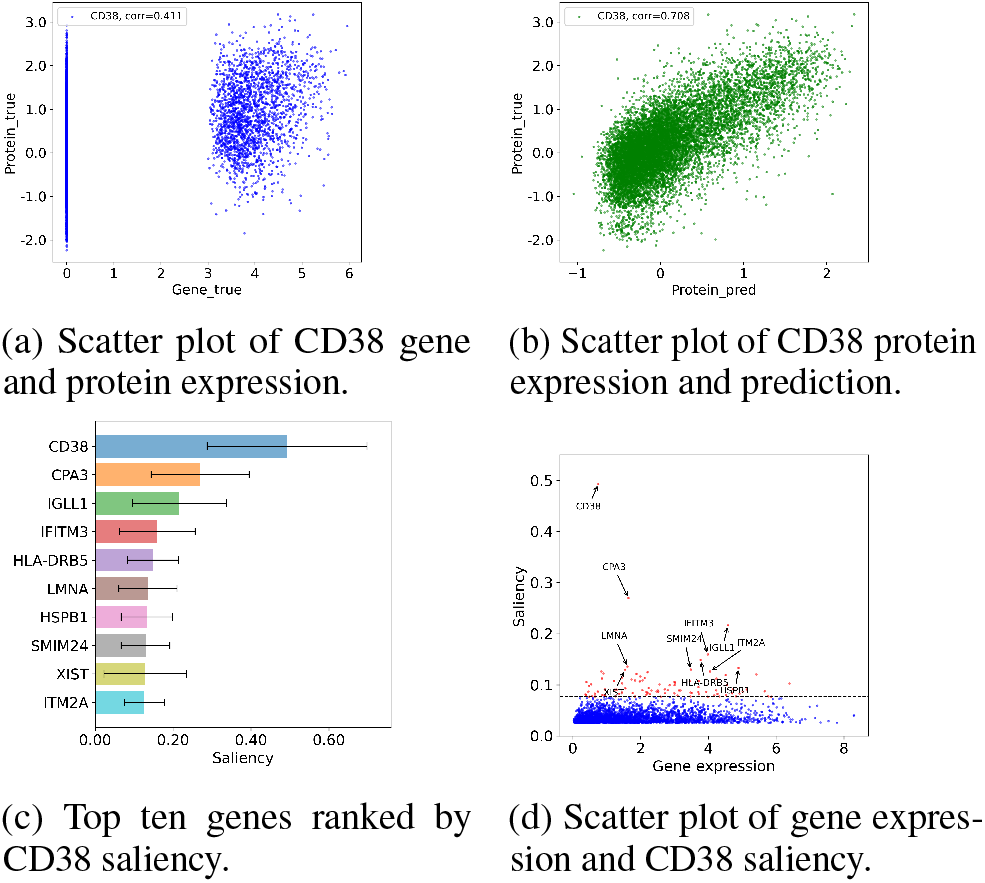
Gene-protein relationship analysis for protein CD38.

Myeloid cells contribute to immunotherapy resistance; however, their role in response to checkpoint inhibition (CPI) in anti-PD-1 refractory cancers is unclear. Next we explore the second CITE-seq data set from (Keenan et al., 2022), where the researchers conclude that CD14+ monocytes linked to anti-PD-1 resistance in human biliary cancer cause T cell paralysis. T cells are dysfunctional when co-cultured with monocytes that express high levels of Tim3.

We first show the expression correlation between surface protein Tim3 and its coding genes *HAVCR2* (0.152, Figure 3a) as well as our model’s prediction (0.649, Figure 3b). Figure S2b depicts the learned time embedding, and the proximity of weeks 2 and 3 indicates that an interpretable embedding has been learned to reveal their inherent similarity. The saliency ranking of the ten most prominent Tim3 genes is depicted in Figure 3c. Due to the intrinsically low correlations between the protein Tim3 and gene *HAVCR2*, the gene has not been ranked highly (Figure 3d) and has had less of an impact on our prediction of Tim3. GSEA ranks *phagosome* as the most significant pathway (Figure 4b), correlating with the finding that PD-1 signaling can polarize macrophages to an M2 phenotype, cause defects in phagocytosis, and impair antitumor immunity (Keenan et al., 2022).

**Figure 3.**
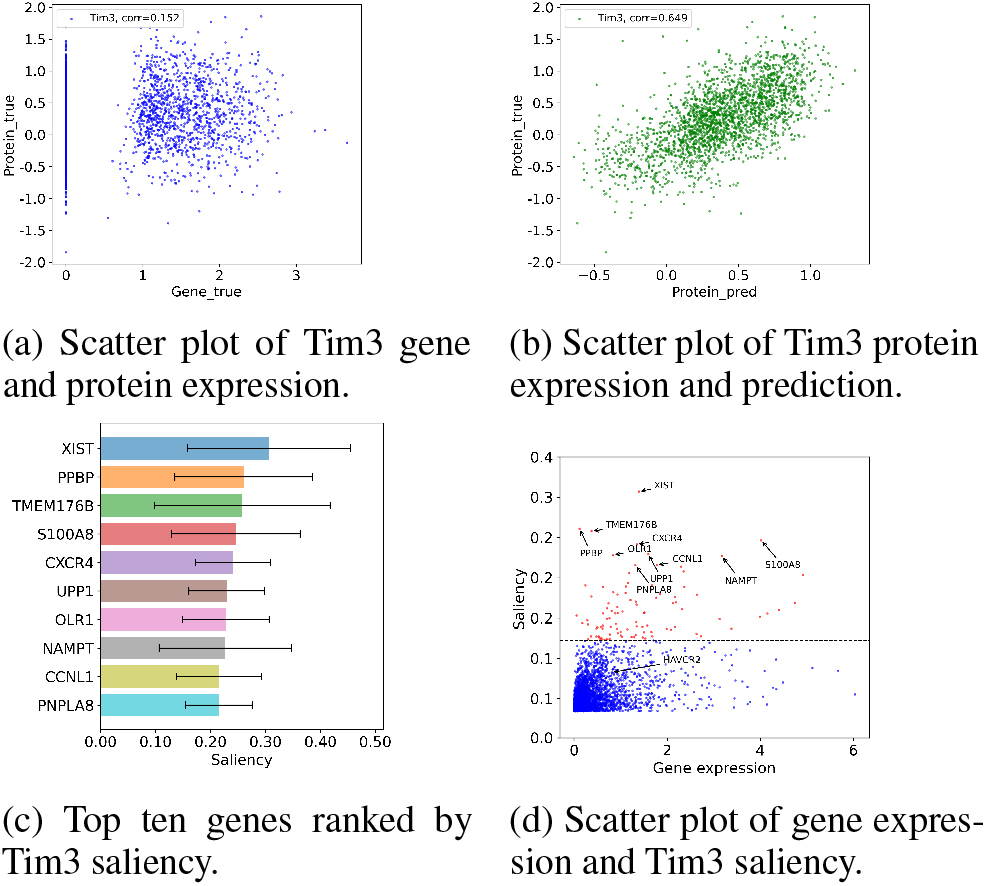
Gene-protein relationship analysis for protein Tim3.

**Figure 4.**
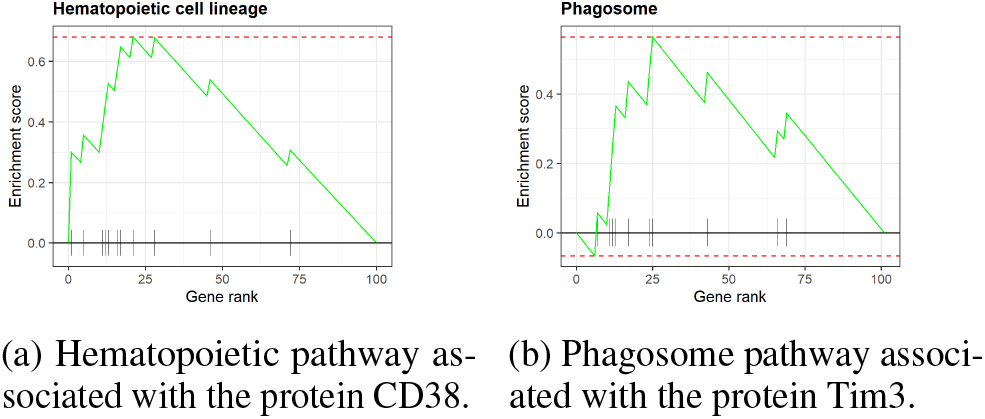
GSEA plots depict the top enriched pathways associated with CD38 and Tim3 proteins. Gene rank represents the position of each gene based on its saliency.

### 4.4. CrossmodalNet is scalable

We simulate three CITE-seq data sets of varying sizes as input and evaluate the scalability of our model by comparing the total training time with other baseline methods. Without using GPUs, our model exhibits a 7.4-to 14.3-fold faster running speed than the average of baseline methods tested on equivalent hardware (Intel Xeon 6248R at 3.0 GHz with 24 GB RAM requested, Table S2). Our model is expected to run faster by enabling GPU implementation. The scalability of our model would allow large experiments with more than thousands of genes.

## 5. Discussion

We present CrossmodalNet as a highly interpretable and scalable model that can be generalized for the prediction of proteomics data from transcriptomics data. Our experiments show that our model with the customized adaptive loss outperforms benchmarking methods including linear, nonlinear and tree-based models, as evidenced by higher Pearson correlation coefficients in most benchmarking scenarios. In practice, once scRNA-seq data of a cell system is available, our model can accurately estimate the patterns of surface proteins. Thus, our model provides an in-silico alternative to CITE-seq experiments and may facilitate the generation of hypotheses and the design of experiments.

Our method for decomposing temporal gene expression into basal and time embeddings of cells is an advancement step towards understanding the mechanisms that govern gene-protein regulation and cell state transitions. The time embedding offers insights into the underlying mechanisms of cell system development and is useful for predicting protein expression in a time-specific manner. In addition, our inference for basal embedding uncovered by our model is biologically interpretable, and we demonstrate that causal gene-protein relationships that provides a fundamental understanding of how genetic information is translated into functional proteins can be deduced from this inference. This quantitative understanding is essential for identifying cellular development and can be expanded to detect disease-causing genes, develop new drugs, and understand complex cellular processes.

Future work will be directed to investigate strategies for incorporating gene-protein prior knowledge into model training, and to extend this work to model scATAC-seq data (Buenrostro et al., 2015). As scATAC-seq and scRNA-seq are naturally causally related, we expect to discover more robust causal relations underlying the central dogma of molecular biology.

## Data availability

The implementation of CrossmodalNet is available at: https://github.com/yjgeno/Multimodal_22

## Acknowledgements

This study was funded by the U.S. Department of Defense (GW200026) for J.J.C. We acknowledge the use of advanced computing resources provided by Texas A&M High Performance Research Computing in conducting parts of this research.

## A. Supplementary tables

**Table S1.** Random search spaces for hyperparameter tuning.

| MODULE | HYPERPARAMETER | DEFAULT VALUE | SEARCH SPACE |
| --- | --- | --- | --- |
| MLP GENERAL | BATCH NORMALIZATION | TRUE | CHOICE([TRUE, FALSE]) |
| MLP GENERAL | LEARNING RATE | 0.001 | QLOGUNIFORM(1E-4, 1E-1, 5E-5) |
| MLP GENERAL | WEIGHT DECAY | 5E-6 | $10^{randint(-3, -7)}$ |
| MLP1 | DROPOUT | 0.05 | CHOICE([0, 0.05, 0.15, 0.3]) |
| MLP1 | LATENT DIMENSION | 512 | CHOICE([256, 512]) |
| DISCRIMINATOR | HIDDEN DIMENSION | 128 | CHOICE([128, 32]) |
| DISCRIMINATOR | REGULARIZATION | 0.5 | QUNIFORM(0, 2, 0.1) |
| DISCRIMINATOR | GRADIENT PENALTY | 0.4 | QUNIFORM(0, 2, 0.1) |
| DISCRIMINATOR | LEARNING RATE | 0.0087 | QLOGUNIFORM(1E-4, 1E-1, 5E-5) |
| DISCRIMINATOR | WEIGHT DECAY | 5E-5 | $10^{randint(-3, -7)}$ |
| DISCRIMINATOR | LEARNING STEP | 3 | CHOICE([3, 5, 10]) |
| GRADNORM | WEIGHT LEARNING RATE | 0.01475 | QLOGUNIFORM(1E-4, 1E-1, 5E-5) |
| GRADNORM | ALPHA | 0.5 | QUNIFORM(0, 3, 0.1) |

**Table S2.** Running time (s) on synthetic data sets (samples, features).

| MODEL | D1 | D2 | D3 |
| --- | --- | --- | --- |
|  | (1,000, 3,000) | (3,000, 5,000) | (10,000, 8,000) |
| CROSSMODALNET | 22.17 | 147.66 | 791.84 |
| MLP | 89.96 | 302.67 | 1604.53 |
| LIGHTGBM | 638.01 | 4364.12 | 29606.51 |
| LINEAR REG. | 48.53 | 850.14 | 23845.28 |
| RIDGE REG. | 11.16 | 97.32 | 1099.67 |
| SCIPENN | 42.28 | 154.48 | 626.36 |

## B. Supplementary figures

**Figure S1.**
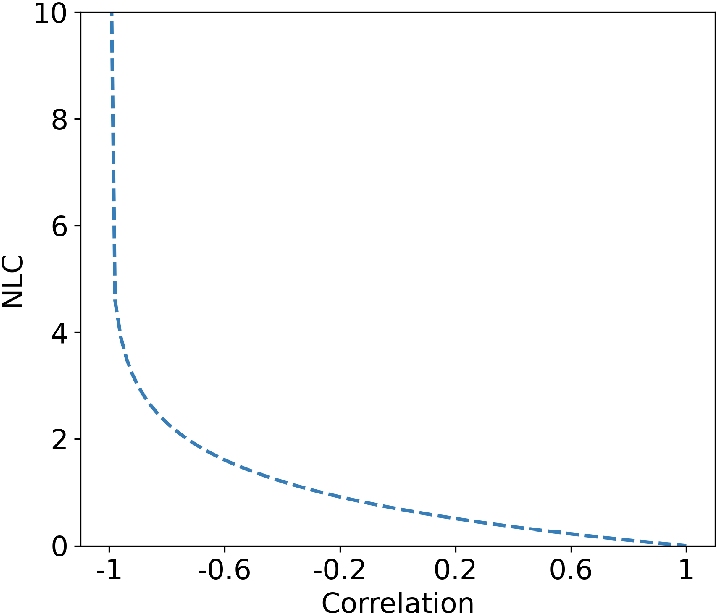
NLC loss curve across correlations.

**Figure S2.**
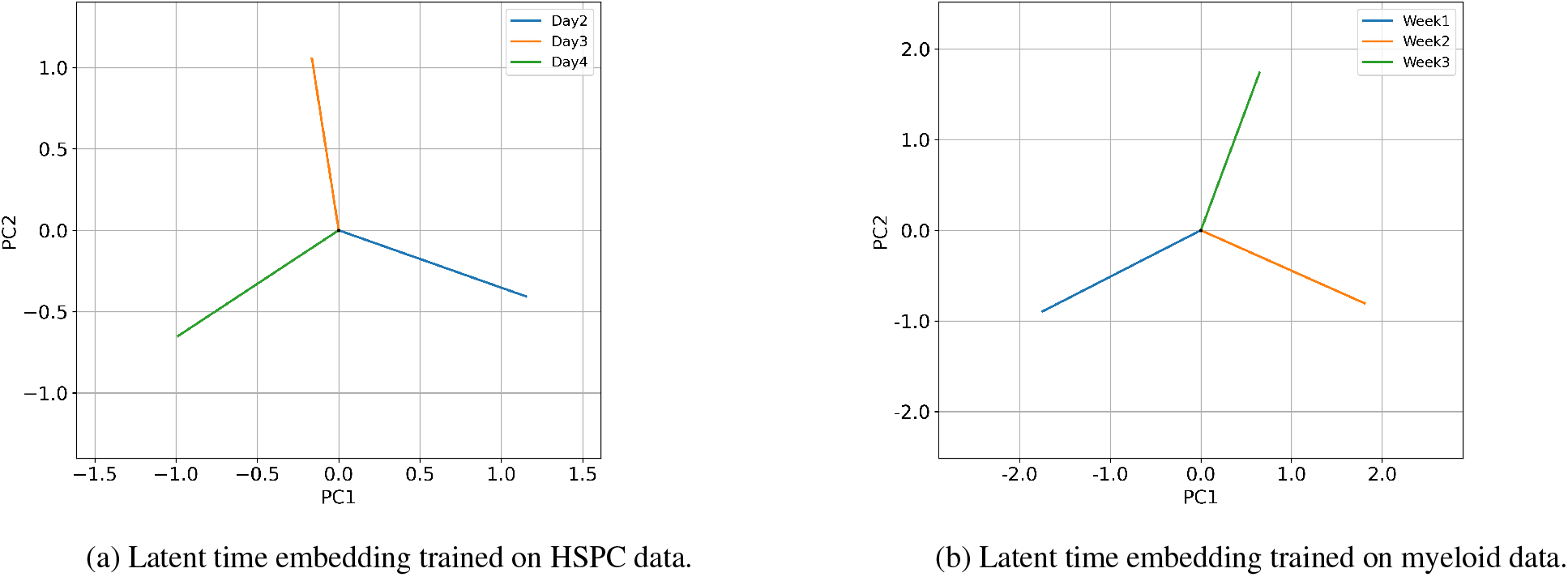
PCA representation of trained latent time embeddings.

## References

Buenrostro, J. D., Wu, B., Litzenburger, U. M., Ruff, D., Gonzales, M. L., Snyder, M. P., Chang, H. Y., and Green-leaf, W. J. Single-cell chromatin accessibility reveals principles of regulatory variation. Nature, 523(7561): 486–490, 2015.

Cellarity et al. Open problems in single-cell analysis. Kaggle. Note: https://www.kaggle.com/competitions/open-problems-multimodal, 2022.

Chen, Z., Badrinarayanan, V., Lee, C.-Y., and Rabinovich, A. Gradnorm: Gradient normalization for adaptive loss balancing in deep multitask networks. In International conference on machine learning, pp. 794–803. PMLR, 2018.

Falcon et al., W. Pytorch lightning. GitHub. Note: https://github.com/PyTorchLightning/pytorch-lightning, 3, 2019.

Gayoso, A., Steier, Z., Lopez, R., Regier, J., Nazor, K. L., Streets, A., and Yosef, N. Joint probabilistic modeling of single-cell multi-omic data with totalvi. Nature methods, 18(3):272–282, 2021.

Han, X., Jiang, Z., Liu, N., and Hu, X. G-mixup: Graph data augmentation for graph classification. In International Conference on Machine Learning, pp. 1–9. PMLR, 2022a.

Han, X., Jiang, Z., Liu, N., Song, Q., Li, J., and Hu, X. Geometric graph representation learning via maximizing rate reduction. In Proceedings of the ACM Web Conference 2022, pp. 1226–1237, 2022b.

Hao, Y., Hao, S., Andersen-Nissen, E., Mauck III, W. M., Zheng, S., Butler, A., Lee, M. J., Wilk, A. J., Darby, C., Zager, M., et al. Integrated analysis of multimodal single-cell data. Cell, 184(13):3573–3587, 2021.

Inecik, K., Uhlmann, A., Lotfollahi, M., and Theis, F. J. Multicpa: Multimodal compositional perturbation autoencoder. bioRxiv, pp. 2022–07, 2022.

Jindal, A., Gupta, P., and Sengupta, D. Discovery of rare cells from voluminous single cell expression data. Nature communications, 9(1):4719, 2018.

Ke, G., Meng, Q., Finley, T., Wang, T., Chen, W., Ma, W., Ye, Q., and Liu, T.-Y. LightGBM: A highly efficient gradient boosting decision tree. Advances in Neural Information Processing Systems, 2017.

Keenan, B. P., McCarthy, E. E., Ilano, A., Yang, H., Zhang, L., Allaire, K., Fan, Z., Li, T., Lee, D. S., Sun, Y., et al. Circulating monocytes associated with anti-pd-1 resistance in human biliary cancer induce t cell paralysis. Cell Reports, 40(12):111384, 2022.

Lakkis, J., Schroeder, A., Su, K., Lee, M. Y., Bashore, A. C., Reilly, M. P., and Li, M. A multi-use deep learning method for cite-seq and single-cell rna-seq data integration with cell surface protein prediction and imputation. Nature Machine Intelligence, pp. 1–13, 2022.

Lample, G., Zeghidour, N., Usunier, N., Bordes, A., Denoyer, L., and Ranzato, M. Fader networks: Manipulating images by sliding attributes. Advances in neural information processing systems, 30, 2017.

Liaw, R., Liang, E., Nishihara, R., Moritz, P., Gonzalez, J. E., and Stoica, I. Tune: A research platform for distributed model selection and training. arXiv preprint arXiv:1807.05118, 2018.

Ling, H., Jiang, Z., Liu, M., Ji, S., and Zou, N. Graph mixup with soft alignments. In International Conference on Machine Learning. PMLR, 2023.

Lotfollahi, M., Klimovskaia Susmelj, A., De Donno, C., Hetzel, L., Ji, Y., Ibarra, I. L., Srivatsan, S. R., Naghipourfar, M., Daza, R. M., Martin, B., et al. Predicting cellular responses to complex perturbations in high-throughput screens. Molecular Systems Biology, pp. e11517, 2023.

Mulè, M. P., Martins, A. J., and Tsang, J. S. Normalizing and denoising protein expression data from droplet-based single cell profiling. Nature communications, 13(1):2099, 2022.

Osorio, D., Zhong, Y., Li, G., Huang, J. Z., and Cai, J. J. sctenifoldnet: a machine learning workflow for constructing and comparing transcriptome-wide gene regulatory networks from single-cell data. Patterns, 1(9):100139, 2020.

Osorio, D., Zhong, Y., Li, G., Xu, Q., Yang, Y., Tian, Y., Chapkin, R. S., Huang, J. Z., and Cai, J. J. sctenifoldknk: An efficient virtual knockout tool for gene function predictions via single-cell gene regulatory network perturbation. Patterns, 3(3):100434, 2022.

Pascanu, R., Mikolov, T., and Bengio, Y. On the difficulty of training recurrent neural networks. In International conference on machine learning, pp. 1310–1318. Pmlr, 2013.

Paszke, A., Gross, S., Massa, F., Lerer, A., Bradbury, J., Chanan, G., Killeen, T., Lin, Z., Gimelshein, N., Antiga, L., Desmaison, A., Kopf, A., Yang, E., DeVito, Z., Raison, M., Tejani, A., Chilamkurthy, S., Steiner, B., Fang, L., Bai, J., and Chintala, S. Pytorch: An imperative style, high-performance deep learning library. In Advances in Neural Information Processing Systems 32, pp. 8024–8035. Curran Associates, Inc., 2019.

Quake, S. R. The cell as a bag of rna. Trends in Genetics, 37(12):1064–1068, 2021.

Restani, P., Beretta, B., Fiocchi, A., Ballabio, C., and Galli, C. L. Cross-reactivity between mammalian proteins. Annals of Allergy, Asthma & Immunology, 89(6):11–15, 2002.

Santurkar, S., Tsipras, D., Ilyas, A., and Madry, A. How does batch normalization help optimization? Advances in neural information processing systems, 31, 2018.

Simonyan, K., Vedaldi, A., and Zisserman, A. Deep inside convolutional networks: Visualising image classification models and saliency maps. arXiv preprint arXiv:1312.6034, 2013.

Stoeckius, M., Hafemeister, C., Stephenson, W., Houck-Loomis, B., Chattopadhyay, P. K., Swerdlow, H., Satija, R., and Smibert, P. Simultaneous epitope and transcriptome measurement in single cells. Nature methods, 14 (9):865–868, 2017.

Velten, L., Haas, S. F., Raffel, S., Blaszkiewicz, S., Islam, S., Hennig, B. P., Hirche, C., Lutz, C., Buss, E. C., Nowak, D., et al. Human haematopoietic stem cell lineage commitment is a continuous process. Nature cell biology, 19 (4):271–281, 2017.

Yang, Y., Li, G., Zhong, Y., Xu, Q., Lin, Y.-T., Roman-Vicharra, C., Chapkin, R. S., and Cai, J. J. sctenifoldxct: A semi-supervised method for predicting cell-cell interactions and mapping cellular communication graphs. Cell Systems, 2023.

